# The SIRAH force field 2.0: Altius, Fortius, Citius

**DOI:** 10.1101/436774

**Authors:** Matías R. Machado, Exequiel E. Barrera, Florencia Klein, Martín Sóñora, Steffano Silva, Sergio Pantano

## Abstract

A new version of the coarse-grained (CG) SIRAH force field for proteins has been developed. Modifications to bonded and non-bonded interactions on the existing molecular topologies significantly ameliorate the structural description and flexibility of a non-redundant set of proteins. The SIRAH 2.0 force field has also been ported to the popular simulation package AMBER, which along with the former implementation in GROMACS expands significantly the potential range of users and performance of this CG force field on CPU/GPU codes.

As a non-trivial example of application, we undertook the structural and dynamical analysis of the most abundant and conserved calcium-binding protein, namely, Calmodulin (CaM). CaM is constituted by two calcium-binding motifs called EF-hands, which in presence of Calcium specifically recognize a cognate peptide by embracing it. CG simulations of CaM bound to four Calcium ions in the presence or absence of a binding peptide (holo and apo forms, respectively), resulted in good and stable ion coordination. The simulation of the holo form starting from an experimental structure sampled near-native conformations, retrieving quasi-atomistic precision. Removing the binding peptide enabled the EF-hands to perform large reciprocal movements, comparable to those observed in NMR structures. On the other hand, the isolated peptide starting from the helical conformation experienced spontaneous unfolding, in agreement with previous experimental data. However, repositioning the peptide in the neighborhood of one EF-hand not only prevented the peptide unfolding but also drove CaM to a fully bound conformation with both EF-hands embracing the cognate peptide, resembling the experimental holo structure.

Therefore, SIRAH 2.0 showed the capacity to handle a number of structurally and dynamically challenging situations including metal ion coordination, unbiased conformational sampling, and specific protein-peptide recognition.

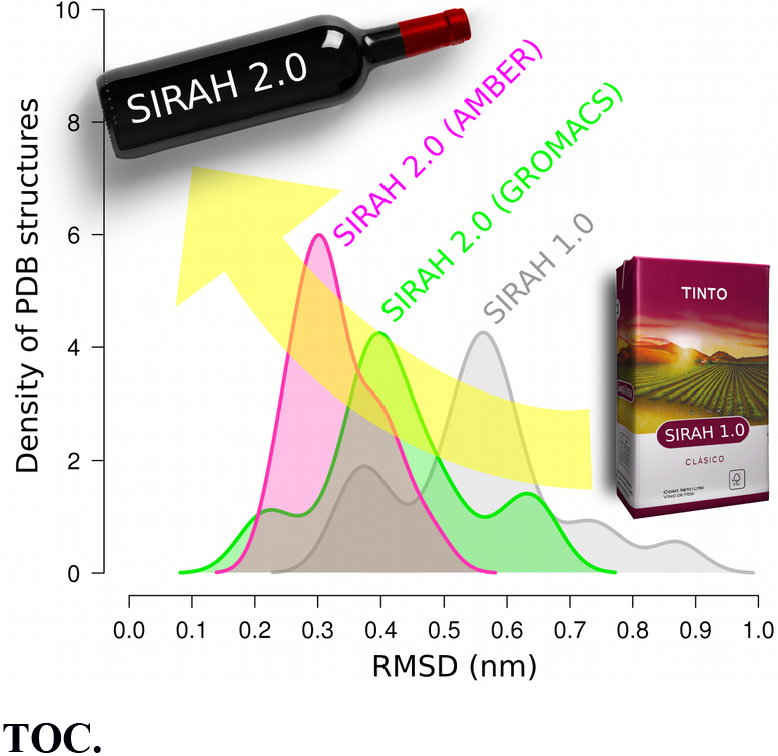
TOC.

## INTRODUCTION

Coarse-grained (CG) models are increasingly gaining popularity because of the possibility to boost the sampling capabilities of Molecular Dynamics (MD) simulations ^1,2^. A large palette of models has been developed following a variety of mapping schemes and strategies for parameter development ^3–11^. Briefly, the CG parameterization can be performed following bottom-up or top-down approaches to fit structural, dynamical o thermodynamic properties. In the first case, frequently referred as systematic derivation, parameters are derived from higher-level simulations (fully atomistic or even quantum). Iterative Boltzmann Inversion, Force Matching or Inverse Montecarlo are among the most popular techniques for bottom-up parameterization ^12,13^. In the second case, parameters are derived aiming to reproduce a selected set of experimental data relevant for the phenomenon under study. These approaches use a variety of fitting techniques and are frequently guided by physicochemical intuition ^1,2^. Regardless the case, the computational speed up, comes at the cost of resigning atomistic resolution ^14,15^. Therefore, continuous efforts are devoted by several groups to improve even well established CG force fields ^5,16–20^. During the last years, our group has developed a CG force field called SIRAH (http://www.sirahff.com). This force field currently contains parameters for proteins ^21^, nucleotides ^22^, aqueous solvent ^23^, and lipids ^24^. Moreover, it is suitable for multiscale simulations ^25–28^ in combination with popular atomistic force fields ^29^. While CG DNA and multiscale simulations showed a very good accuracy ^25,30,31^, in particular cases, the protein representation still shows relatively large deviations from experimental structures. Indeed, simulation of a set of non-redundant protein-DNA complexes revealed that the protein rather than the DNA experienced significantly larger deviations from the initial structure ^32^. Therefore, we sought to improve the protein’s parameters to obtain a better structural representation and facilitate the implementation of the force field in popular MD packages. Here we present a significant update of our force field (SIRAH 2.0), which includes: i) a series of modifications to bonded and non-bonded parameters of the amino acids, while keeping the same topologies; ii) description of neutral states for Aspartate and Glutamate residues; iii) improved mapping compatibility from different force fields and experimental structures; iv) updated functionalities to the set of analysis and visualization tools (SIRAH Tools ^33^); v) porting to AMBER; vi) the possibility to profit from GPU acceleration in AMBER and GROMACS codes.

Improving the structural description of amino acids resulted in a decrease of the averaged RMSD on a protein data set by nearly 0.1 nm in relation to the previous version. The capabilities of SIRAH 2.0 were also evident from the study of different states and components of the Calmodulin (CaM) system, on which we obtained an unprecedented qualitative and fully unbiased description of the protein-peptide recognition.

## METHODS

### Re-parameterization of the Model

SIRAH uses a classical Hamiltonian common to most all-atoms potentials to describe particle-particle interactions. Briefly, bonds and angles were described by harmonic terms (e.g.: *k*_b_/2*(*r*-*r*_b_)^2^, with force constant *k*_b_ and equilibrium values *r*_b_), while Fourier expansions were used for dihedrals. Non-bonded contributions were accounted by Coulomb and Particle Mesh Ewald summation (PME ^34,35^), while Lennard-Jones (LJ) terms were calculated using the standard 12-6 expression (i.e.: 4* *ɛ*\*[(*σ*/*r*)^12^-(*σ*/*r*)^6^]). Lorentz-Berthelot combination rules were used for most atom-types pairs. However, specific LJ parameters were set for tuning interactions between some bead pairs (see below). The AMBER scaling factors SCEE=1.2 and SCNB=2.0 were used for the 1-4 non-bonded interactions.

Aimed to preserve compatibility within the existing force field, we kept the same amino acid’s topologies and introduced a series of modifications in bonded and non-bonded parameters. For the optimization procedure we used the following two-steps top-down approach:

1. <u>Testing:</u> a given modification was tested for structural stability on a suitable system. For instance, secondary structure modifications were tested on all α or all β structures, disulfide bridges were tested on a Cysteine rich protein family called Defensin, Calcium coordination was tested on EF-hands and C2 domains, etc.
2. <u>Validation:</u> Changes were accepted if they improved a set of structural descriptors averaged on a set of proteins used to test the robustness of the first version of SIRAH ^21^. The validation set contained 18 structures known to be monomeric, shorter than 120 amino acids, with one replica in the asymmetric unit, without ligands and containing α and/or β secondary structure elements. The structural descriptors used were Root Mean Square Deviation (RMSD), Solvent Accessible Surface (SAS), Radius of Gyration (RGYR), and native contacts. Using state of the art GPU accelerated desktop computers, the simulation of the entire set of proteins for time windows of 1 μs was accomplished within a couple of days.

In the next sections we describe the changes made to the SIRAH force field.

#### Uniform masses

Starting from this new version of SIRAH, all beads in the force field have been assigned a mass of 50 a.u., except for the recently developed supra CG water model ^28^.

#### Improved backbone torsionals

The CG beads composing the protein backbone are placed in the positions of the real atoms N, O and Cα (Figure 1A). This rather detailed description allowed a simple and direct geometrical equivalence between all-atoms (AA) and CG dihedral angles. It also granted an unbiased representation of different secondary structure elements at CG level. This mapping scheme remained unchanged. However, in the original version of the force field (SIRAH 1.0) ^21^, the CG potentials for Ψ_CG_ 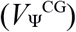 and Ф_CG_ 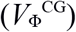 were implemented as a sum of three and four periodic terms, respectively. This representation was substituted by Fourier expansions containing two and nine terms, respectively (Table 1). Despite simplifying the functional form of 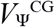, its energy landscape remained essentially unchanged (Figure 1B). On the other hand, we increased the complexity of 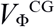 to allow for two separated minima at parallel and antiparallel β-sheet conformations. Energy barriers for some transitions were also reduced to increase the peptide’s flexibility. It can be deduced from Figure 1B that this new combination of 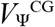 and 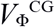 functions generated minima compatible with the canonical secondary structure regions of the Ramachandran plot. Another major limitation in the previous force field version was the impossibility to describe peptide bonds in *cis* conformation. This was solved by adding a minimum at 0º to the torsional angle function of Ω_CG_ 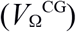, accounting for the possibility to adopt *cis* and *trans* peptide bonds (Table 1, Figure 1B).

**Figure 1.**
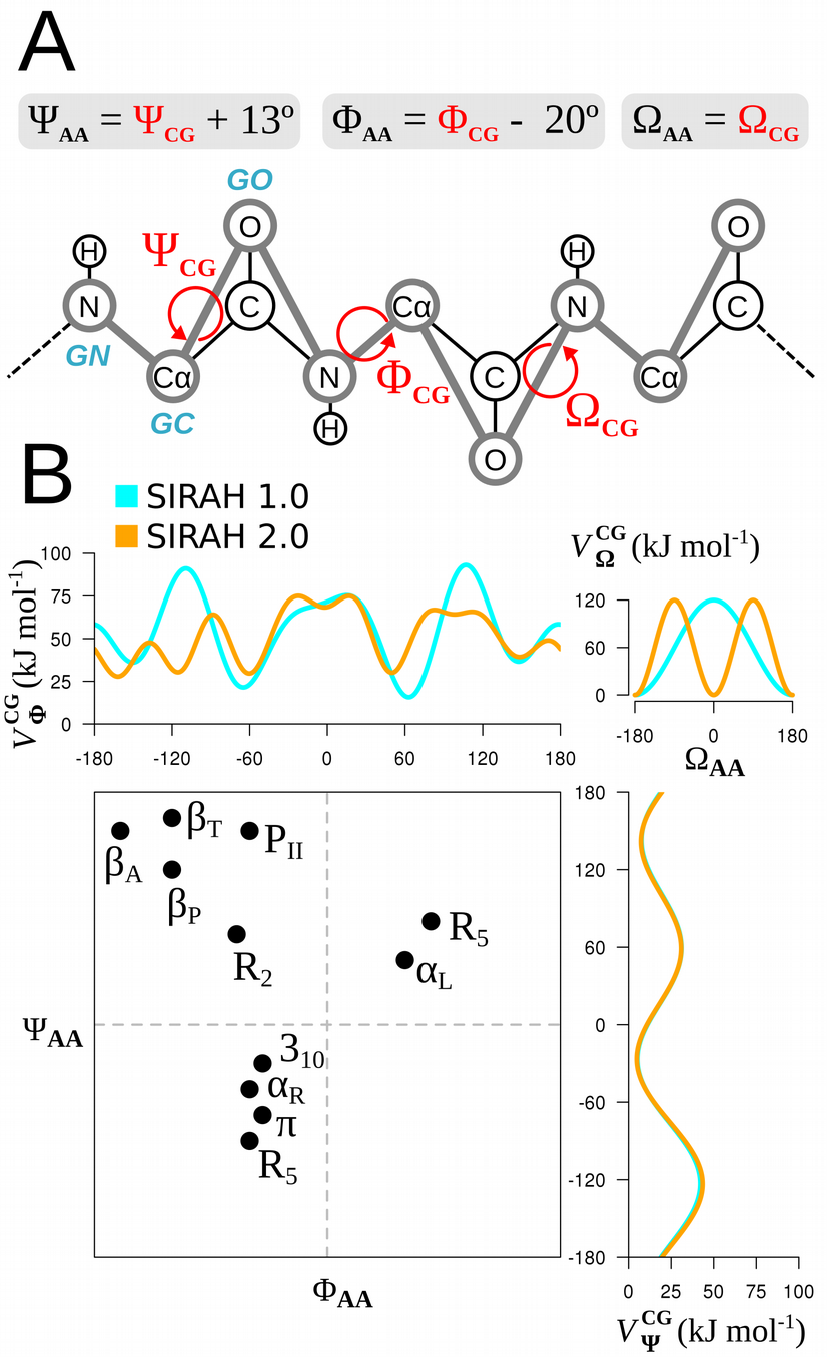
Backbone definition and Ramachandran plot. A) Mapping and geometrical equivalence between AA and CG torsional angles along the backbone. CG beads are named in blue italic characters and their connectivity is indicated as thick gray lines. B) Graphic representation of CG dihedral potential. To facilitate the comparison against canonical secondary structure elements in the atomistic Ramachandran plot, the torsional energy landscape is plotted using the AA geometrical space, which simply implies shifting the CG functions according to the transformations indicated on panel A (e.g. 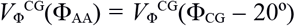). The position of right-handed α-helix (α_R_), 3_10_-helix (3_10_), π-helix (π), left-handed α-helix (α_L_), 2-fold ribbon (R_2_), 5-membered ring (R_5_), polyproline-II (P_II_), parallel β-sheet (β_P_), antiparallel β-sheet (β_A_) and twisted β-sheet (β_T_) were taken from ref ^36^.

**Table 1.** New torsional parameters in SIRAH 2.0 for proteins.^a^

| **Table 1.** New torsional parameters in SIRAH 2.0 for proteins.<sup>a</sup> |  |  |  |  |
| --- | --- | --- | --- | --- |
| Dihedral angle | m | $k_m$ (kJ mol<sup>-1</sup>) | $n_m$ | $\alpha_{0m}$ (deg) |
| $\Psi_{CG}$ (GN<sub>i</sub>-GC<sub>i</sub>-GO<sub>i</sub>-GN<sub>i+1</sub>) | 1 | 6.4 | 1 | -146.5 |
|  | 2 | 15.4 | 2 | 89.4 |
| $\Phi_{CG}$ (GO<sub>i-1</sub>-GN<sub>i</sub>-GC<sub>i</sub>-GO<sub>i</sub>) | 1 | 11.6 | 1 | 49.0 |
|  | 2 | 4.9 | 2 | -38.8 |
|  | 3 | 9.4 | 3 | 42.5 |
|  | 4 | 8.2 | 4 | 79.8 |
|  | 5 | 0.4 | 5 | -114.4 |
|  | 6 | 5.2 | 6 | -52.0 |
|  | 7 | 8.9 | 7 | -87.8 |
|  | 8 | 1.5 | 8 | 103.1 |
|  | 9 | 1.3 | 9 | 65.9 |
| $\Omega_{CG}$ (GC<sub>i-1</sub>-GO<sub>i</sub>-GN<sub>i</sub>-GC<sub>i</sub>) | 1 | 60.0 | 2 | 180.0 |
| <sup>a</sup> The torsional potential is expressed according to GROMACS' definition: $V = \sum_m k_m [1 + \cos(n_m \alpha - \alpha_{0m})]$. | | | | |

#### Increased side chain flexibility

Although the amino acids’ topologies did not change (Figure 2A), some angular force constants were reduced to allow for a wider range of side chain motion. Specifically, the force constant at the side chains of Arginine and Lysine (angles defined by the beads GC-BCG-BCZ and GC-BCG-BCE, Figure 2A) were changed from 10.0 to 2.0 kJ mol^-1^ rad^-2^. Similarly, the force constant at the side chains of Serine, Threonine and reduced Cysteine (angles defined by beads GC-BOG-BPG and GC-BSG-BPG) were changed from 50.0 to 10.0 kJ mol^-1^ rad^-2^.

**Figure 2.**
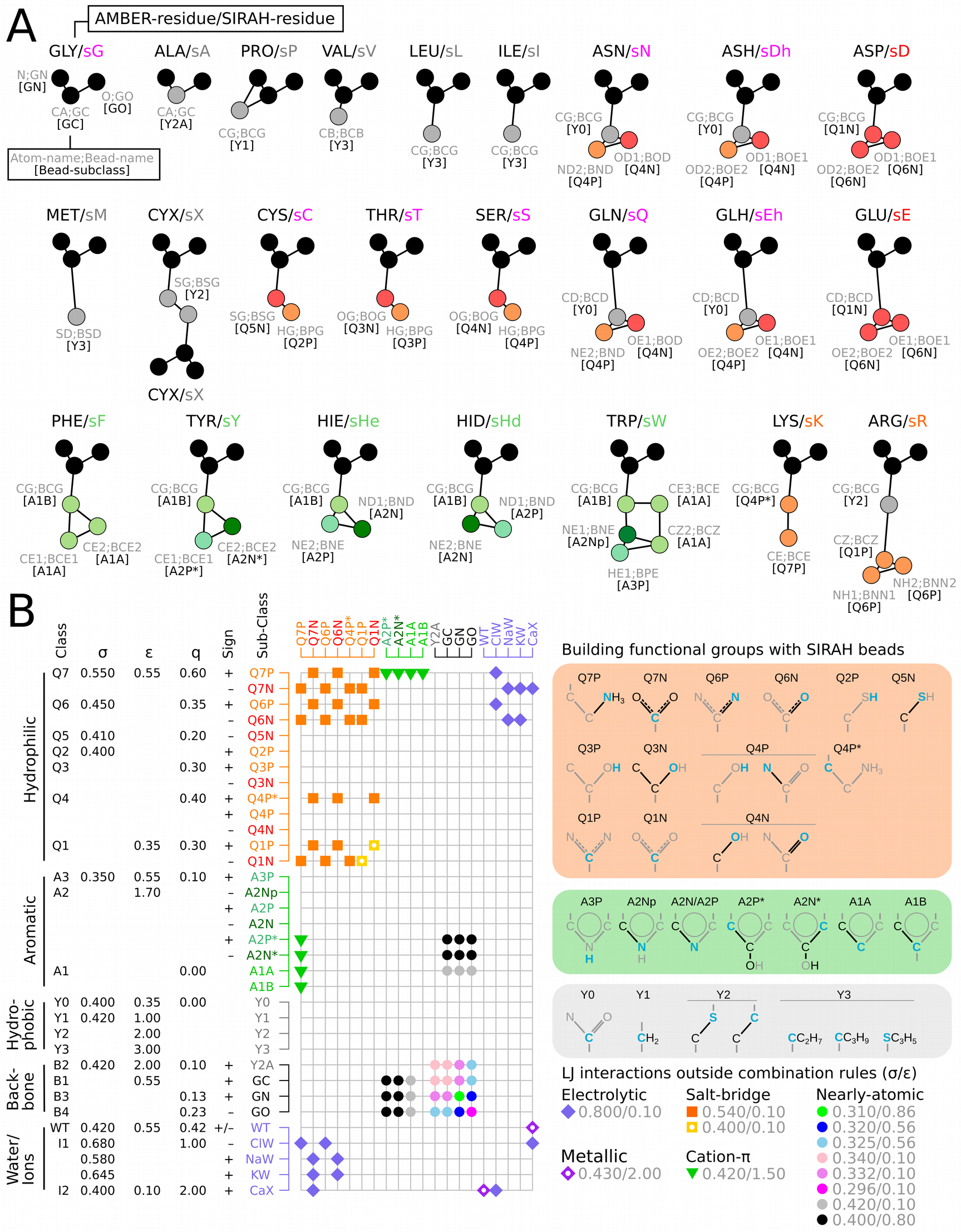
Comprehensive view of SIRAH 2.0 for proteins. A) Topologies of CG amino acids. CG beads are placed in the position of real atoms, colors indicate their physicochemical character as defined in panel B. B) Bead-classes, sub-classes and non-bonded parameters in SIRAH 2.0. The LJ interaction matrix is shown for beads having interactions outside the Lorentz-Berthelot combination rules as defined by the corresponding symbol in the grid. Right panels rationalize the SIRAH beads in terms of chemical functional groups. Mapped atoms are in blue, while represented fragments also embrace atoms and bonds in black.

#### Improved non-bonded interactions

It is worth recalling that the protein model was developed for explicit solvent simulations in combination with the CG water model called WatFour (WT4 for shortness) ^23^, which is composed by four interconnected beads in a tetrahedral arrangement, each carrying a partial charge of +/- 0.41 e and σ, ɛ LJ parameters of 0.42 nm and 0.55 kJ mol^-1^, respectively (Figure 2B). In this work we introduced a series of modifications to the non-bonded parameters of several beads aimed to improve the hydrophobic/hydrophilic balance at side chains and backbone. Beads were divided in five groups (Hydrophilic, Aromatic, Hydrophobic, Backbone and Water/Ions) and within each group different classes were defined according to their charge module and LJ parameters (Figure 2B). Within each class we distinguished subclasses, which could be characterized by the charge sign and/or a special interaction (see next paragraph). This newly introduced classification was somehow related to specific functional or chemical characteristics. For example, the polar group in Serine was represented by two charged particles of the same bead-class with opposing charge (i.e. different bead-subclass). Table 2 summarizes all modifications incorporated to non-bonded parameters.

**Table 2.**
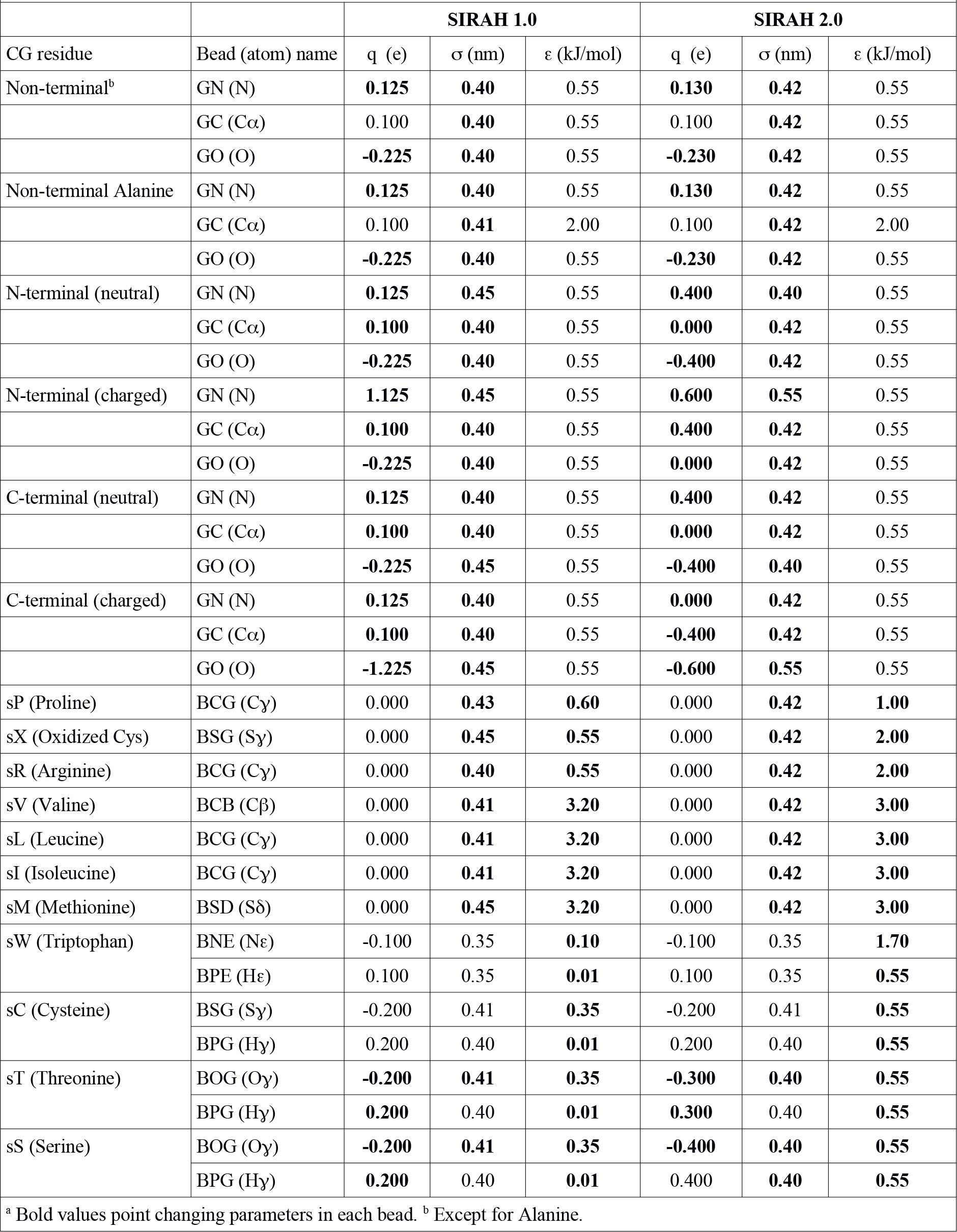
List of all changes in non-bonded parameters from SIRAH version 1.0 to 2.0.^a^

#### Interactions outside Lorentz-Berthelot combination rules

When using simplified models, one single set of non-bonded parameters per bead may not be general enough to capture diverse chemical specificities. Setting differential LJ interactions outside the Lorentz-Berthelot combination rules between some bead-subclasses provided a convenient workaround for this issue. In particular, we defined the following special interaction types: i) Nearly-atomic; ii) Salt-bridge; iii) Cation-π; iv) Electrolytic and v) Metallic.

i. Nearly-atomic: In this case, LJ interactions were tuned to permit closer contacts between specific beads. In particular, these allow for intra-backbone Hydrogen bond-like contacts and the necessary atom packing in α-helices and β-sheets. In SIRAH 2.0, these interactions were also used to modulate contacts between the backbone and aromatic side chains. This also improved the representation of Hydrogen bonds frequently present between buried Tyrosine and Tryptophan and backbone atoms.
ii. Specific salt-bridge interactions were kept from the original version. These were set to compensate the excess of electrostatic attraction between opposite charged amino acids.
iii. Cation-π interactions were added to account for the special interaction between aromatic moieties and cationic groups.
iv. Electrolytic interactions occurred among ions in solution or electrolytes and charged residues. All LJ interactions in such cases were set to σ 0.80 nm and ɛ 0.10 kJ mol^-1^ to avoid spurious formation of ionic pairs. In this sense, it could be important to recall that simple electrolytes in the SIRAH force field were represented with van der Waals (vdW) radii corresponding to their second solvation shell ^23^. Hence, they were meant to modulate the ionic strength in the solution and not specific ion-protein interactions.
v. Metal ions pose significant challenges to CG and even to classical force fields. However, the use of differential LJ interactions with protein, solvent and anionic electrolytes (Cl^-^, see Figure 2) allowed us to achieve a rather accurate description of Ca^2+^ bound to CaM. Although Ca^2+^ constitutes the only current example in the force field, preliminary studies indicate that similar strategies can be applied to other metallic ions. This issue will be addressed in deeper detail in a forthcoming publication.

A comprehensive view of the bead-types, non-bonded parameters, specific interactions and chemical functional groups represented by the CG moieties is presented in Figure 2B.

#### Improved terminal residues

The possibility to represent neutral or zwitterionc termini was preserved in SIRAH 2.0 but the non-bonded characteristics of the terminal beads were changed and the charge was distributed between the N/C terminal beads and their corresponding Cα as indicated in Table 2.

### SIRAH in AMBER

Since SIRAH was implemented with the same Hamiltonian function used by common atomistic force fields, it is portable to virtually any MD engine. Indeed, the DNA parameters for implicit or explicit solvent simulations at CG ^22,23^ or dual-resolution (AA/CG)^25^ were originally developed within the AMBER suite. In this work we implemented the entire new version of SIRAH on AMBER. To ensure transferability, the new dihedral potentials 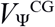, 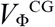 and 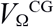 were defined using sets of functions without repeating the same periodicity (Table 1). In addition, improper torsions were implemented to preserve their original definition in Darré et al. 2015 ^21^ and also to account for the way in which the AMBER’s Leap module writes them into the topology. Regardless their definition in the parameter file, Leap only requires the central atom to be given at third position while the rest is re-sorted in alphabetical order by atom-type except for wild cards, which are placed first. Hence, the amino acid “L” chirality was defined by the improper torsion X-GN-GC-GO (where “X” is an AMBER wild card standing for any bead-type), with an energy barrier of 100 kJ mol^-1^, a periodicity of 1 and a phase of - 35.0 degrees. All LJ interactions outside the combination rules (Figure 2B) were set in the LJEDIT section of the parameter file, which is a feature available since AMBER 14 ^37^. Importantly, running SIRAH 2.0 on AMBER GPU codes ^38,39^ requires versions 14 or later, since older ones don’t read the full LJ information from the topology.

### SIRAH Tools

We made a big effort to provide an easy to use, plug and play force field. In that sense, we developed a set of scripts (SIRAH Tools) to help the mapping, backmapping, visualization and analysis of SIRAH trajectories ^33^. Several improvements were made for the new force field version, including more generic mapping files, updated selection macros and vdW radii for the VMD plugin ^40^. The updated SIRAH package for AMBER and GROMACS, including tutorials, is freely available at http://www.sirahff.com.

#### Mapping atomistic systems to SIRAH

It is possible to map experimental or simulated all-atom structures to SIRAH as long as they contain the mapping atoms. Disulfide bonds as well as different protonation states for Histidine, Glutamic and Aspartic acid residues could be automatically recognized (see Figure 2A). We used the AMBER residue and atom denominations for best mapping compatibility. We now added support for naming schemes of CHARMM, GROMOS, GROMACS and OPLS force fields. However, some ambiguity will always remain in the residue naming, especially regarding protonation states, which may lead to a wrong mapping. For example, SIRAH Tools maps by default the residue name “HIS” to a Histidine protonated in Nε regardless the actual protonation state. If protonation on Nδ is required, the residue name in the PDB file has to be edited from HIS to HID (or HSD in CHARMM). Similarly, protonated Glutamic and Aspartic acid residues must be named “GLH” and “ASH”, otherwise they will be treated as negative charged residues. These kind of situations need to be carefully checked by the users. In all cases the residues preserved their identity upon mapping and backmapping the structures.

### Computational details

#### Validation set

Initial structures were taken from the PDB and processed with the PDB2PQR server ^41^ to add protons at pH 7 and possible missing atoms. The structures were then mapped to CG using SIRAH Tools ^33^. The CG models were solvated using pre-equilibrated WT4 molecules in octahedral boxes of 2.0 nm size from the solute (see *Practical notes on system solvation*). An ionic strength of 0.15 M was set by randomly replacing WT4 molecules by Na^+^ and Cl^-^ CG ions.

The simulation protocol consisted on: 1) Solvent and side chains relaxation by 5000 steps of energy minimization, imposing positional restrains of 1000 kJ mol^-1^ nm^-2^ on backbone beads corresponding to the nitrogen and carbonylic oxygen (named GN and GO, respectively, Figure 1A); 2) Full system relaxation by 5000 steps of unrestrained energy minimization; 3) Solvent equilibration by 5 ns of MD in NVT ensemble at 300 K imposing positional restrains of 1000 kJ mol^-1^ nm^-2^ on the whole protein; 4) Protein relaxation by 25 ns of MD in NVT ensemble at 300 K imposing positional restrains of 100 kJ mol^-1^ nm^-2^ on GN and GO beads; 5) Production simulation in NPT ensemble at 300 K and 1 bar. Non-bonded interactions were treated with a 1.2 nm cutoff and PME for long-range electrostatics. A timestep of 20 fs was used in MD simulations. Snapshots were recorded every 100 ps for analysis.

It is worth noting that the use of a standard classical Hamiltonian with a 12-6 term for LJ interactions within a generally smoothed CG landscape and in the absence of topological constraints makes the force field susceptible to close contacts. In such cases gentler minimization/equilibration protocols can sensibly ameliorate the structural stability.

Simulations were run using GROMACS 2018.2 (http://www.gromacs.org) and/or AMBER 16 (http://ambermd.org) GPU codes ^38,39,42^. Owing to different features, algorithms and implementations in both MD software packages, specific flags were set in each case. In GROMACS, PME and neighbor searching were computed each 10 integration steps, considering a Verlet cutoff-scheme of 1.4 nm. Automatic tuning of these options was not allowed by setting the verlet-buffer-drift flag to −1. Solvent and solute were coupled separately to V-rescale thermostats ^43^ with coupling times of 2 ps. The system’s pressure was controlled by the Parrinello-Rahman barostat ^44,45^ with a coupling time of 8 ps. On the other hand, in AMBER, PME was calculated every integration step owing to code restrictions, and the neighbor list was updated whenever any atom had moved more than 1/2 a non-bonded “skin” of 0.2 nm. The whole system was coupled to the Langevin thermostat ^46^ with a collision frequency of 50 ps^-1^ and to the Berendsen’s barostat ^47^ with a relaxation time of 1 ps. The results of different simulations showed to be rather robust to the choice of thermostat, barostats and their coupling parameters.

#### Practical notes on system solvation

The placement of solvent molecules around the solute is an important step in the system setup. The most common strategy consists on filling the computational box with clusters of pre-equilibrated solvent molecules and then removing the overlapping ones. Although fast, this strategy leaves intermolecular interstices, which are eventually fixed in subsequent simulation steps. In CG systems this issue might become more pronounced due to the higher granularity of the particles. In practice, using the actual bead’s sizes of SIRAH produced poorly solvated proteins, no solvent molecules were present up to 0.5 nm away. Such condition had dramatic effects during the energy minimization and equilibration as charged amino acids at the surface of the protein virtually interacted “in vacuum” during initial steps leading to important structural distortions. A way to improve the initial solute hydration was allowing for small overlaps with solvent molecules when generating the solvation box. In AMBER this was accomplished by scaling the solvent-solute interaction radii by 0.7 when adding the WT4 molecules. In GROMACS, the solvation was done using the default radii of 0.105 nm for atoms not present in the vdW database (vdwradii.dat) and then removing the WT4 molecules within 0.3 nm from the solute. In all cases, eventual clashes were relaxed during the solute-restrained energy minimization.

#### Calmodulin systems

CaM simulations were performed using as starting conformer the first model in the NMR family deposited under the PDB id: 2KNE ^48^, which was reported as the most representative in the ensemble. The holo complex contained CaM bound to four Calcium ions and a cognate peptide from the Plasma Membrane Calcium ATPase (PMCA). The apo form of CaM was generated by removing the binding peptide from the holo complex. The CaM-peptide recognition was analyzed by taking the conformer at time = 1 μs from the CaM apo simulation and manually placing the PMCA peptide in the proximity of the C-terminal EF-hand. The docking procedure only implied orienting Tryptophan 1093 of PMCA towards the hydrophobic patch which belongs to the protein-peptide interface and avoiding steric clashes. This arbitrary conformation was used to start an unbiased MD simulation. The simulation of the isolated peptide was started using the same conformation present in the structure 2KNE, which featured 81% helical content. A control peptide of sequence YSEEEERRRR was started in fully helical (canonical) conformation. All simulations were performed using the same setup and protocol previously described for the Validation set.

### Calculated properties

RMSD and RGYR were calculated on Cα atoms, in the former case only residues at α-helices or β-sheets in the experimental structure were considered. The secondary structure content was calculated using SIRAH Tools ^33^. Proteins’ solvent accessible surfaces (SAS) were calculated with GROMACS’ utility g_sas by setting a probe radius of 0.21 nm (i.e, the radius of a WT4 bead) and by defining a zero charge for hydrophobic beads. The vdW radii corresponding to SIRAH 2.0 beads were used. Protein contacts were calculated using a 0.8 nm cutoff distance between Cα atoms. Native contacts were defined on the experimental structure and its conservation was measured as the retained percentage along the trajectory. The contacts’ accuracy was defined as the ratio between conserved native contacts and the total occurring contacts. All measured properties on the validation set were calculated at CG level from the last 100 ns of simulation.

## RESULTS

### Validation of SIRAH 2.0

The modifications described in the previous section were validated against a set of proteins using both AMBER 16 and GROMACS 2018.2 packages. Both sets of results were compared against the same set of proteins used to test the robustness of SIRAH 1.0 ^21^. Simulations were performed in triplicate for each individual protein and results summarized in Figure 3A. The implementations on AMBER and GROMACS showed slightly different results, which can be ascribed to the different algorithms and approximations implemented in each MD engine. Nevertheless, for both cases we obtained a reduction on the average value of different structural descriptors with respect to the original version. The relative error measured by comparing the average deviations form the initial (experimental) structure indicated relatively small improvements in RGYR and the conservation of native contacts (Figure 3A). However, the accuracy of such contacts showed a more sensible decrease in relation to SIRAH 1.0. A more sizable progress was found for the SAS, suggesting a better balance between hydrophobic and hydrophilic surfaces. The detailed list of results obtained for each protein in the benchmark data set is reported in Supplementary Table ST1.

**Figure 3.**
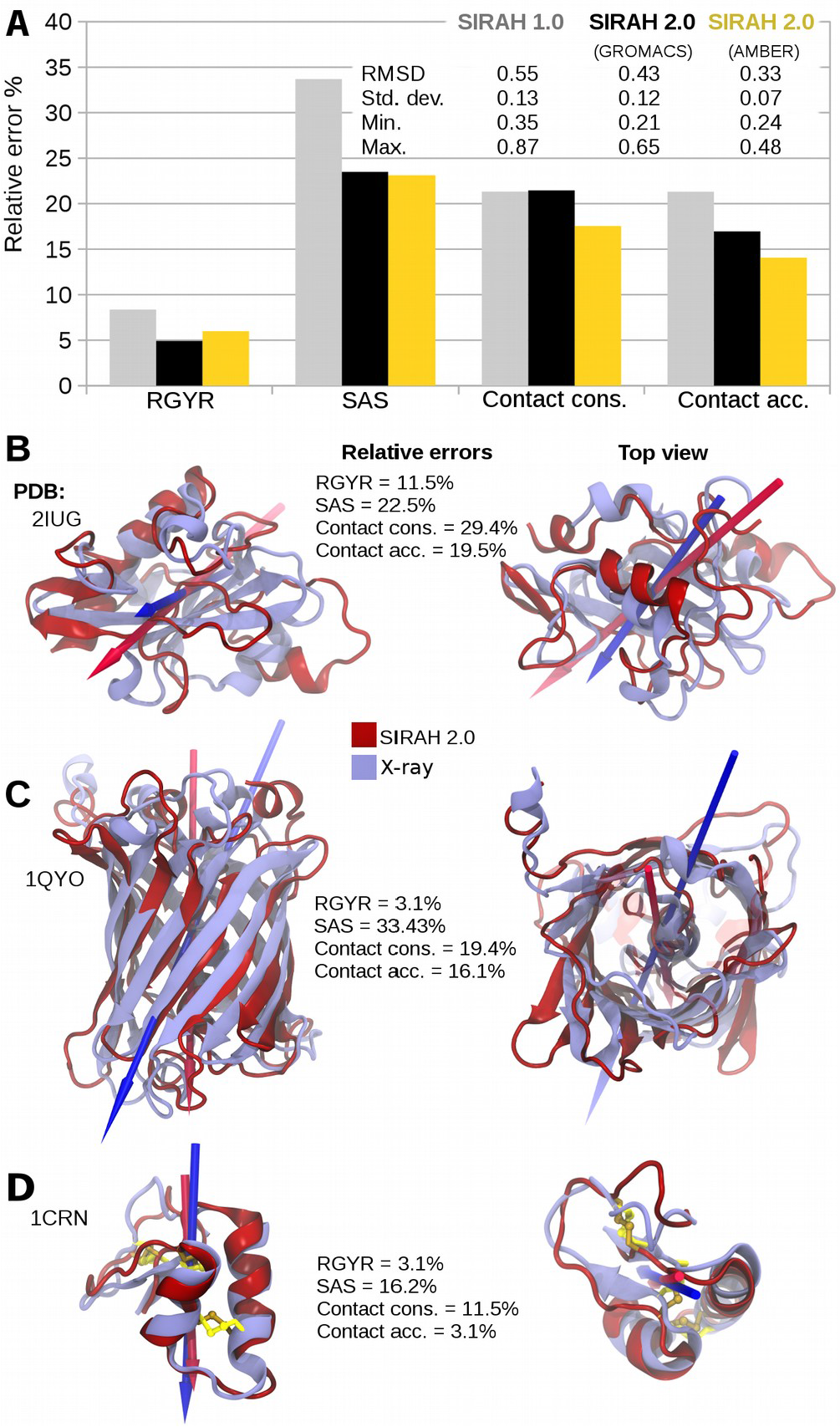
Comparison of structural descriptors among different SIRAH versions and implementations. A) Averaged relative error on structural descriptors in reference to experimental structures. The values of SIRAH 1.0 were taken from Darré et al. 2015 ^21^. B-D) Structural alignment of last frames from worst, average and best performing proteins, respectively. Arrows indicate the dipole moments calculated using the AMBER charge distribution on crystallographic and backmapped structures. Disulfide bridges present in structure 1CNR and kept during the CG simulation are shown in yellow.

The quadratic nature of the RMSD made it worth to take a closer look as a descriptor of structure quality. For the validation set, we obtained a reduction of more than 0.1 nm in the average RMSD values in relation to the old force field version regardless the MD engine (Figure 3A). Worthy, not only the average was reduced but also the minimum and maximum values registered a consistent decrease, suggesting a higher robustness to handle different structural scaffolds. As a gauge of the quality of the structural reproduction that can be expected from SIRAH 2.0 we briefly analyze the results of three simulations obtained from the worst, average and best performing proteins (PDB ids 2IUG ^49^, 1QYO ^50^, and 1CRN ^51^, respectively, Figures 3B-D). Aimed to facilitate the comparison we backmapped the last conformers of each simulation using SIRAH Tools and superimposed them on the corresponding experimental structures.

The worst performing protein, 2IUG, corresponds to the SH2 amino terminal domain of the Pi 3-Kinase P85. It presents a α+β topology and belongs to the SCOP family d.93.1. Protein 2UIG features as main secondary structure elements a central three-stranded antiparallel β-sheet encased by two helical segments. Additionally, the structure is decorated with three small helical and β-sheet elements (Figure 3B). Superimposing the last conformer after 1 μs of simulation performed with GROMACS against the experimental structure evidenced distortions according to a final RMSD value of 0.65 nm with a low conservation of native contacts. Despite this relatively large value, some secondary structure elements were still recognizable and errors in structural descriptors as RGYR and SAS remained within the error observed for the whole set of proteins (Figure 3A and supplementary Table ST1). Hence, the protein maintained its degree of compactness and the correct burial of the hydrophobic core.

The three replicates of protein 1QYO simulated using the AMBER engine presented a RMSD deviation of 0.31 nm, which nearly matched the average of the distribution (Figure 3A and supplementary figure 1). This protein corresponded to an engineered variant of the Green Fluorescent Protein (GFP), which preserved the folding without creating a chromophore inside ^50^. It featured the typical central helix surrounded by 11 antiparallel β-strands, which was globally well maintained (Figure 3C).

The best RMSD performing protein was Crambin, identified with the PDB id 1CRN. Crambin is a small protein from the family of thionins, which act as toxins for most mammalian cells. It has been extensively studied by a series of techniques because of its small size and good structural resolution. It contains three disulfide bridges and two small helices and β-strands. As it can be seen from Figure 3D, the structural comparison was very good, featuring a RMDS of only 0.21 nm. The most evident difference observed between the simulation and the X-ray structure consisted in a slightly higher separation between the two β-strands. It is worth noting that the structural fidelity of this protein registered a significant improvement in going from SIRAH 1.0 to SIRAH 2.0 (Supplementary Figure S1).

Regardless the case, the comparison between the dipole moments calculated on the initial and final conformers evidenced a good degree of colinearity. Hence suggesting that, even in cases of poor structural reproduction as that obtained from protein 2IUG, long-range interactions exerted by the proteins would be well preserved at the CG level.

Taken as a whole, the new version of the force field showed a significant improvement in the structural reproduction of a non-redundant set of proteins. While this constituted an important achievement, it would also be desirable to explore near native conformational states in a completely unbiased manner. This issue is addressed in the following section for a non-trivial test case.

### Exploring the dynamics of Calmodulin

CaM is a calcium-binding protein conserved in all vertebrates and ubiquitously expressed in all Eukaryota, highlighting the biological relevance of this macromolecule. CaM is constituted by two Ca^2+^ binding modules named EF-hands, which are linked by a flexible region. EF-hands are by far the most abundant Ca^2+^ binding domains in all organisms, regulating a plethora of biological processes ^52^. Upon Ca^2+^ binding, the EF-hands expose hydrophobic patches rich in Methionine residues conferring CaM the ability to interact with different targets ^52^. Among others, Ca^2+^ binding to CaM enables the recognition of regulatory peptides present at the Plasma Membrane Calcium ATPases (PMCA). Different isoforms of PMCA are responsible for the Ca^2+^ extrusion from the cytosol or its regulation within cellular microdomains ^53^. In that sense, CaM constitutes an extremely challenging system, as it requires an accurate description of the conformational behavior of a single peptide, Ca^2+^ binding to EF-hands, allosteric movements, large protein flexibility and specific protein-peptide recognition. As a test case, we used the NMR structure of CaM in complex with Ca^2+^ and the regulatory peptide of PMCA (PDB id: 2KNE ^48^), and studied the dynamics of all the components separately and as a holo complex. Furthermore, the previous version of SIRAH was recently used to gain insights onto the ataxia-related G1107D mutation on PMCA ^54^, providing a good comparison between the performance of both versions of our force field.

#### CaM binding peptide

The capability of describing sequence dependent conformational dynamics in oligo-peptides was a celebrated feature of SIRAH 1.0 ^21^. Hence, aimed to dissect the conformational behavior of each component in the CaM holo complex we first focused on the CaM binding peptide. The peptide comprised amino acids 1086 to 1113 of PMCA4. When bound to CaM it presents an 81% of helical content. However, circular dichroism on an isolated PMCA peptide showed negligible amounts of secondary structure ^54^. In close agreement with these observations, our CG simulation of the isolated peptide showed a spontaneous decrease in secondary structure, which started from the terminal residues and then propagated to the center of the molecule (Supplementary Movie 1). As seen from Figure 4A, the helical content decreased in a step-wise fashion, exhibiting metastable steps. Within each step we observed partial refolding events, which were evident from the non-monotonic decrease of the helical content. The helical content of the isolated peptide varied as much as 20% within each step, indicating that up to 4 amino acids left and re-entered into the helical region, underlying the capacity of SIRAH 2.0 to sample a wide range of conformations. The simulation was stopped after 110 μs since only a marginal helical segment remained at the central region.

**Figure 4.**
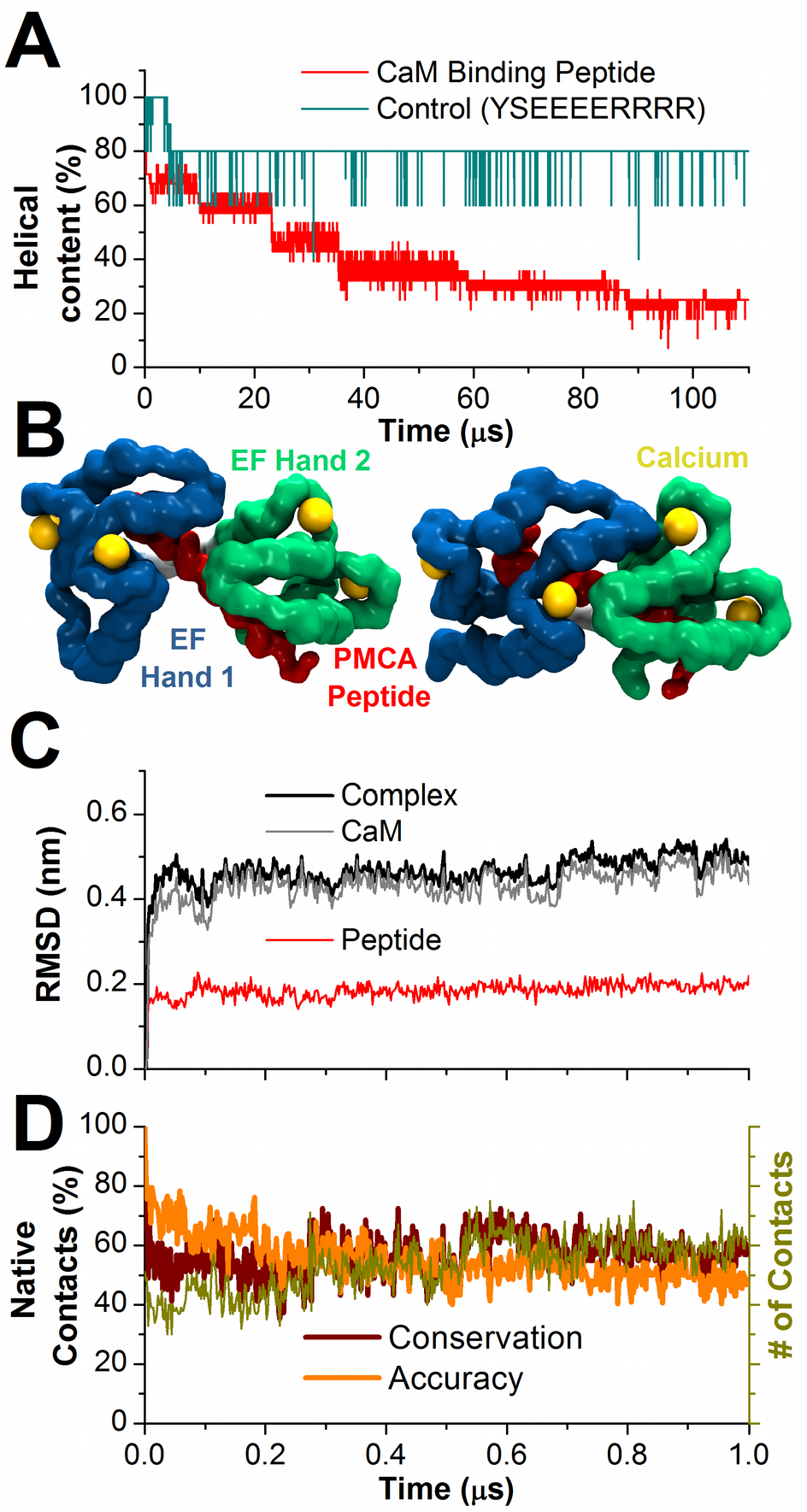
Dynamics of the CaM binding peptide and CaM holo complex. A) Helical content of isolated peptides. B) Initial (left) and final (right) conformations of the CaM holo complex. Only solvent accessible surfaces of the backbone are shown for clarity. C) RMSD traces of the holo complex and its individual constituents. D) Contacts at the CaM-peptide interface. The total number of contacts and percentage of native contacts’ conservation and accuracy are shown in green, dark red and orange traces, respectively.

As a control, we performed a simulation of the peptide YSEEEERRRR, which was shown to retain a highly helical character despite its short extension ^55^. This simulation was started with all amino acids in α helical conformation. In stark contrast to the previous case, this peptide showed a high stability retaining nearly 80% of its helical content (Figure 4A, Supplementary Movie 1). The average helical content over 110 μs was 79%, which is comparable to the approximate 70% determined by circular dichroism experiments and other theoretical predictions ^55^. The helical stability of this peptide putatively originated from the formation of electrostatic interactions between Arginines and Glutamates in a helical context. This arrangement brings together positive and negative moieties placed at positions *i* and *i*+*4*. In agreement with this observation, only the first two amino acids (Tyrosine and Serine) adopted a disordered conformation after 5 μs while all the others leaved the helical conformation in a fleeting fashion (see Supplementary Movie 1).

The comparison between both simulations underlined the fact that, besides the roughness of the topological description, SIRAH 2.0 preserved sequence specificity and unbiased secondary structure flexibility.

#### CaM holo complex

As a second step, we sought to simulate the CaM holo complex as solved by NMR (i.e., including 4 Ca^2+^ ions and the regulatory peptide). Comparison between the initial and final conformers (Figure 4B), suggested a globally good structural conservation. As seen from Figure 4C, the CG simulation produced a stable trajectory reaching a RMSD plateau at nearly 0.5 nm if calculated on all Cα beads of the CaM holo complex (Figure. 4C). This value decreases to 0.45 nm if calculated on the secondary structure elements. This can be considered as a good structural reproduction within the coarseness of the approach. Indeed, a similar simulation performed with SIRAH 1.0 resulted in larger RMSD deviations (about 0.6 nm) ^54^. Yet, it was enough to provide a rationale for the reduced affinity of the Glycine to Aspartate mutation in a CaM binding domain of PMCA, which was not evident from atomistic simulations. Hence, we can expect for SIRAH 2.0 to harbor potential to effectively enhance the understanding of molecular systems at an affordable computational cost.

Calculation of RMSD on CaM and the PMCA peptide separately indicated that the deviation from the initial conformation could be mostly ascribed to CaM, which followed closely the behavior of the entire complex (Figure 4C). In contrast to the simulation of the isolated binding peptide, the presence of the protein context sufficed to stabilize the helical structure of the peptide, which preserved an average helical content of 72%.

Calculation of native contacts’ conservation at the protein-peptide interface rapidly dropped to nearly 60% after minimization and the accuracy of such contacts followed a similar behavior (Figure 4D). This result was consistent with previous simulations on different protein-protein complexes using SIRAH 1.0, where we also observed a similar trend ^21^. In both cases, the loss of native contacts putatively resulted from the intrinsic loss of specificity at CG level. Nevertheless, calculation of the total number of contacts remained nearly constant, experiencing a slight increase with sizable oscillations (Figure 4D). The average number of inter Cα contacts was 58 over the last 100 ns, which is comparable to the initial value of 51, indicating that the protein-protein interface was maintained despite the loss in specificity.

A stringent control for this system was the coordination of Ca^2+^ ions. There are nearly 10000 structures containing Ca^2+^ bound currently reported on the PDB. Hence, a good reproduction of Ca^2+^ coordination must be considered as an important goal for any biomolecular force field. Aimed to define a cut-off distance for the Ca^2+^ coordination we used the GSP4PDB server (https://structuralbio.utalca.cl/gsp4pdb) ^56^ to generate a distance distribution of any amino acid around Calcium ions in all structures reported at the PDB. From the distribution shown in Figure 5A it resulted evident that the first shell of coordinating amino acids was comprised within 0.35 nm. Within this distance, 55% of the amino acids were Aspartate or Glutamate followed by Asparagine (10%) and Glycine (9%), while each of the other residues contributed less than 4%. Aspartate, Glutamate and Asparagine coordinated the metal ion through contacts with oxygen atoms from carboxylate or amide groups in their side chains, while other amino acids interacted via their backbone’s carbonylic oxygen. Hence, we calculated the cumulative radial distribution function of oxygen atoms belonging to the three different moieties (carboxylates, amides and backbone’s carbonyls) around Calcium during the simulation. These distributions were compared against the same quantities measured over the four binding sites and all the 20 conformers in the NMR family of CaM (Figure 5B).

**Figure 5.**
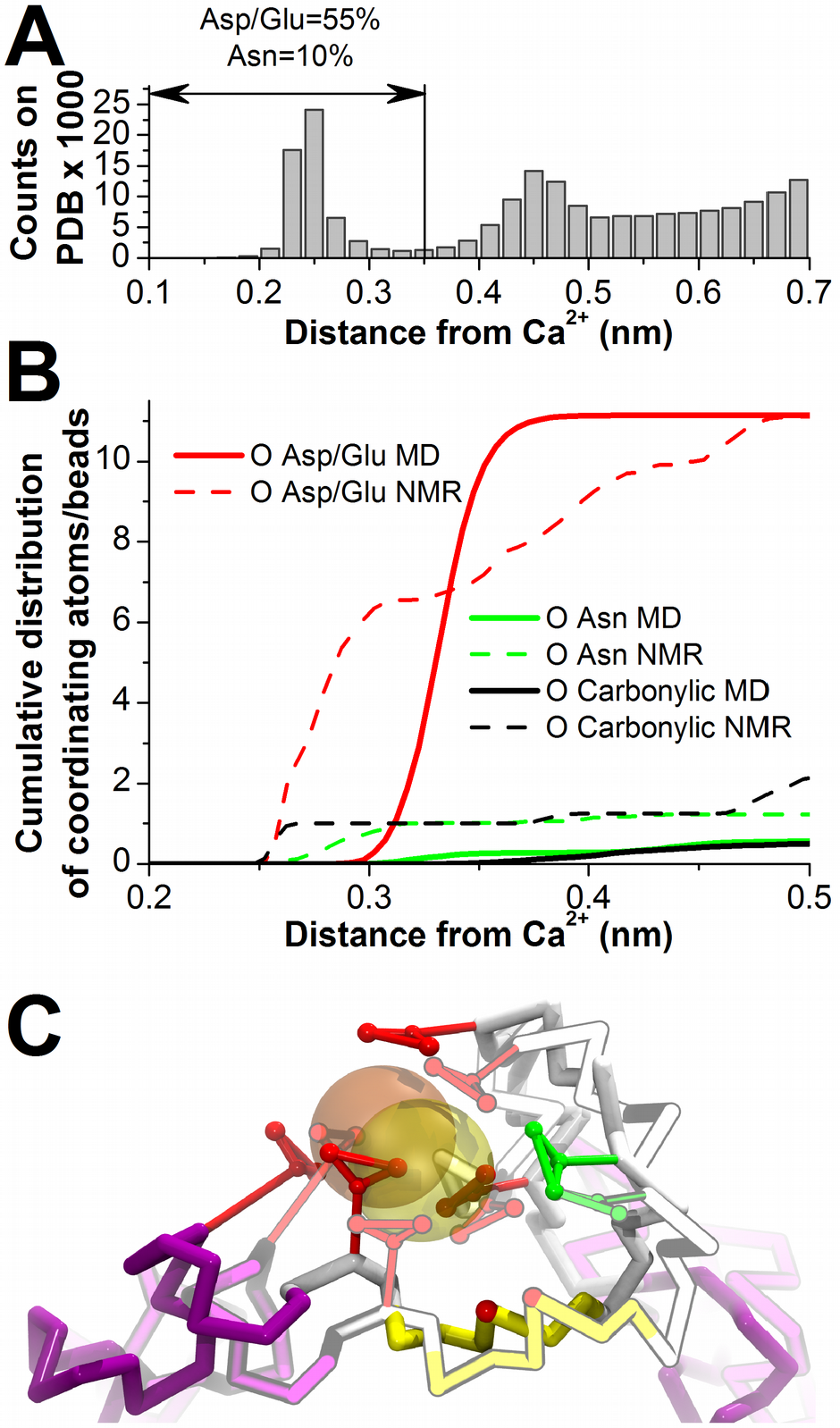
Calcium coordination. A) Radial distribution of amino acids coordinating Ca^2+^ in all reported structures at PDB as calculated from https://structuralbio.utalca.cl/gsp4pdb. B) Cumulative radial distribution functions of oxygen atoms coordinating Ca^2+^ at the binding sites of CaM in the NMR structure 2KNE and the CG simulation. C) Comparison between initial (semi-transparent) and final (opaque) conformations of a Calcium binding site in the simulation of the CaM holo complex. The backbone is colored by secondary structure (α-helix: purple; β-sheet: yellow and coil: white) while side chains of Asp/Glu and Asn are shown in red and green, respectively. A carbonylic oxygen bead participating in the coordination is also shown in red. The initial Ca^2+^ position is colored in yellow.

The NMR and CG cumulative radial distributions of carboxylates converged to the same value within 0.5 nm (red traces in Figure 5B), meaning the overall coordination shell was equally described. However, the higher conformational flexibility available to the atomistic description resulted in a slightly more complex distribution, as compared with the sigmoid-like curve obtained in the CG simulation (Figure 5B). In particular, the larger size of the CG beads in comparison to the atomistic situation produced a right-shift of about 0.1 nm in the carboxylate distribution. A similar feature was observed for amide and carbonyl oxygen. Despite that, the superposition of the initial and final conformers of the Calcium binding site showed a very good structural conservation (Figure 5C). We recently obtained similar results on a membrane channel using SIRAH 1.0 ^57^.

#### CaM apo protein

To study the CaM apo protein we removed the binding peptide from the experimental structure 2KNE of the holo complex and performed a CG simulation with SIRAH 2.0. In contrast to the holo form, the absence of the binding peptide induced RMSD excursions of CaM beyond 2 nm from the bound conformation (compare gray traces in Figures 4C and 6A). As noticed from the RMSD of individual domains, the structural changes were characterized by large separations between both EF-hands, which moved as rather rigid linked objects. Although each EF-hand module also suffered from structural distortions, the force field provided a qualitatively correct description of the exposure of hydrophobic patches in presence of Calcium, in agreement with recent comprehensive analysis of CaM dynamics ^58^. Indeed, both EF-hands spontaneously opened up like a “bear trap” exposing Methionine residues, which were originally buried constituting the protein-peptide interface (see molecular representations of Figure 6A). Therefore, SIRAH 2.0 captured the correct conformational transition at qualitative level.

**Figure 6.**
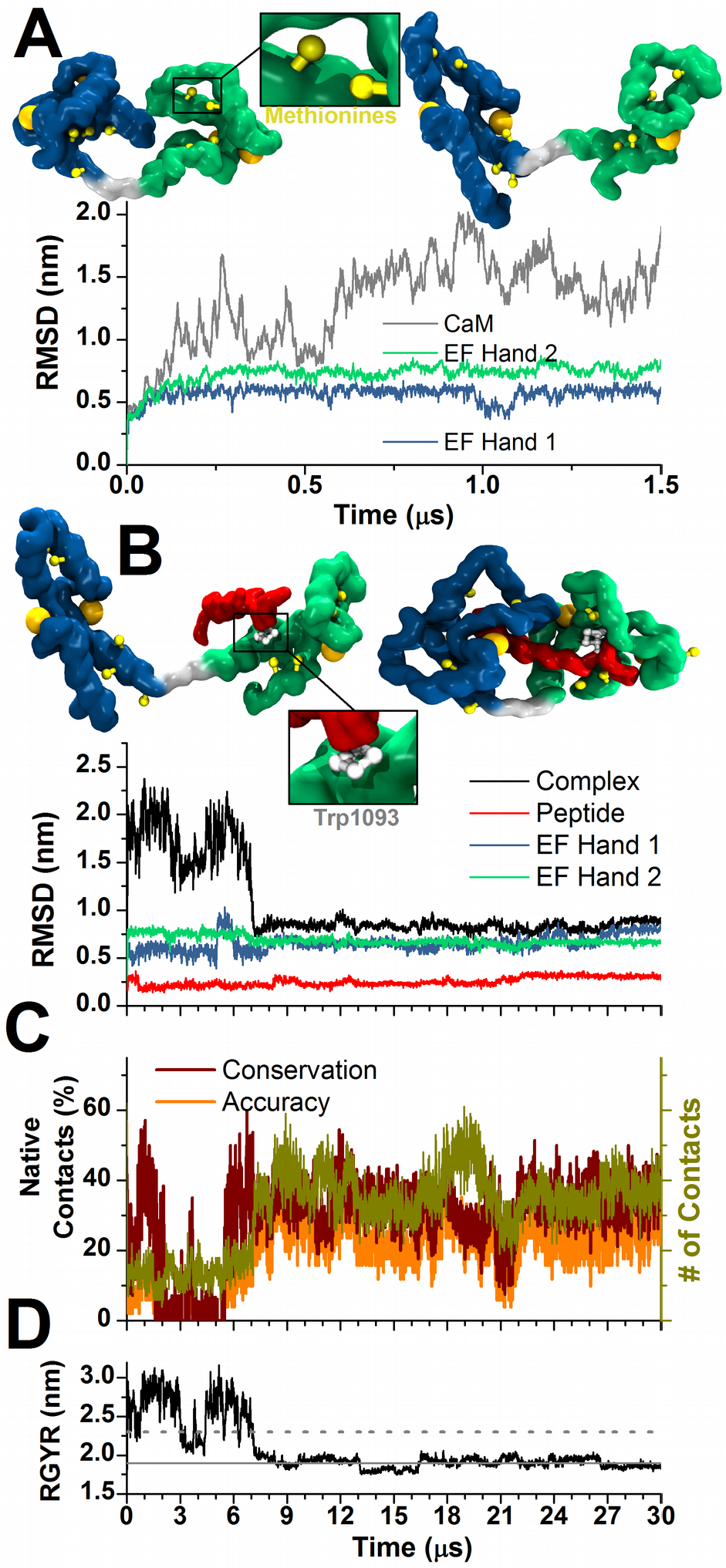
CaM apo protein and PMCA peptide recognition. A) Dynamics of CaM apo after removing the PMCA peptide from the experimental structure. Molecular drawings on top illustrate initial and final conformers. B) PMCA peptide recognition. On top, molecular representations of initial and final structures after manual docking of PMCA peptide to CaM apo and simulation. At bottom, RMSD traces of different parts of the complex taking as reference structure 2KNE. C) Conservation of native contacts (dark red) and accuracy (orange) from structure 2KNE. The number of total contacts is reported in green. D) RGYR of CaM along the simulation. Solid and dashed lines indicate average values calculated on the NMR conformers of structures 2KNE and 1CFF respectively.

#### Calcium free CaM apo

To test the limits of SIRAH 2.0 we sought to investigate CaM apo in its Calcium free form. To this goal we removed all four Ca^2+^ ions and the binding peptide from the experimental structure of the complex and simulated only free CaM apo in aqueous solution.

Already at the equilibration phase it became obvious that the EF-hands in absence of Ca^2+^ underwent severe structural distortions (RMSD over 0.8 nm in each EF-hand domain). This problem arose as a consequence of the strong repulsion between acidic residues coordinating the cations, which were not properly screened by the bulky CG solvent. Indeed, the minimum distance between carboxylate moieties in the NMR structure is nearly 0.5 nm, i.e. barely enough to accommodate a CG water bead, which vdW diameter is 0.42 nm (see Figure 2B). This is likely a common problem to any CG model working with explicit electrostatics and using solvent models representing more than one water molecule in one single bead. A possible work around for this problem could be to perform a detailed analysis of the protonation state of the acidic amino acids in absence of Calcium and introduce eventually protonated Aspartic/Glutamic acids as ASH/GLH in the initial set up. This alternative has shown to be effective to stabilize the conformation of proteins with protonated acidic residues in the original version of the force field ^21^. However, since CaM includes up to 15 protonable residues, we considered that a detailed analysis of possible protonation states goes beyond the scope of the present communication and was not addressed. This could be considered a limitation for our force field, which should to be kept in mind by potential users.

#### Recognition of PMCA peptide by CaM apo protein

Finally, we attempted to mimic the recognition of the PMCA peptide by CaM. To this aim we took a frame from the simulation of the CaM apo protein and manually placed the C-terminal half of the PMCA peptide in the proximity of the second EF-hand domain. The only applied criterion in the docking was facing the Tryptophan 1093 towards the EF-hand (Figure 6A), since this residue has been shown to be crucial for binding ^48^. This complex was then re-solvated and simulated under identical conditions to previous systems. This complex roughly mimicked the PDB structure 1CFF ^59^, in which a short helical peptide is bound to full length CaM making contacts only with its C-terminal EF-hand. It is important to note some clear differences between both conformational states: i) in our case the EF-hand was not wrapped around the peptide; ii) the simulated peptide contained the entire binding peptide; iii) the distances between EF-hands in the simulated and reference complex was significantly different. Nevertheless, since both complexes contain the N-terminal EF-hand free and the C-terminal bound, we used structure 1CFF to lately compare the gross conformational determinants (i.e. RGYR) explored by the NMR structure and our MD simulation.

Calculation of the RMSD on the entire complex immediately identified two well separated conformational states illustrated in Figure 6B. During the first 7 μs CaM showed a dynamics alike to that observed in the apo structure with RMSD reaching more than 2 nm in relation to the holo complex (Figure 6B). Despite this large mobility, the peptide remained associated to the C-terminal EF-hand in a rather nonspecific fashion. After 7 μs, the large unbiased exploration of the conformational space allowed for a random encounter between N-terminal EF-hand in the open “bear trap” conformation, which recognizes the hydrophobic surface of the peptide and wrapped around it. As it can be inferred from the steep decrease in RMSD the recognition happened within the sub μs timescale. After that point the structural descriptors remained stable and dynamics was interrupted after 30 μs. Although the global conformation experienced a significant shift towards the experimental structure of the holo complex, it remained relatively far from it or its CG simulation (compare Figures 4C and 6B). Indeed the RMSD of both EF-hands remained rather insensible when CaM apo embraced the peptide (Figure 6B). This suggests that although the qualitative description of the peptide recognition is correctly reproduced in a fully unbiased way, the atomistic details might be difficult to retrieve within the explored time scale.

To further characterize the bound state found during the simulation we considered additional structural descriptors, namely, contacts at CaM-peptide interface and RGYR of the complex. Evaluation of the contacts, taking those of the experimental complex as native, indicated that during the first 7 μs the percentage of conservation arrived to virtually zero to then rise up to 40% with an accuracy of 30% (Figure 6C), which has to be compared with the 60% obtained in the simulation of the experimental structure of the complex (Figure 4D). Similarly, the total number of contacts raised from 10 at the docking conformation up to 40 at the end of the simulation, which was close to the 51 calculated on the NMR complex.

Finally, RGYR of the complex along the trajectory confirmed the spontaneous evolution of the complex toward a stable compact conformation (Figures 6B and D). In agreement with other descriptors, RGYR showed two clearly separated states. We observed that before the wrapping process took place, the RGYR of the complex experienced large excursions. This behavior could be roughly compared with the RGYR measured on the family of NMR conformers of structure 1CFF (Figure 6D). Indeed, averaging the RGYR on the first 7 μs time window we obtained a value of 2.6 nm, in contrast to the average value of 2.3 nm obtained from the NMR conformers of 1CFF. Remarkably, after CaM completely embraced the PMCA peptide, the RGYR converged to the same quantity measured on the native bound state of the 2KNE structure. Therefore, although the fine structural details of the protein-peptide were not completely retrieved, SIRAH 2.0 produced a very good quantitative description of the unbiased intermolecular recognition.

## CONCLUSIONS

Fully atomistic simulations are well established and constitute a reliable alternative to experimental methods in acquiring detailed structural information, although they suffer from the drawback of an often-prohibitive computational cost. On the other hand, the speed up granted by CG approaches has extended the capability of computational studies to larger size and/or temporal scales, bridging the gap between the available computer power and molecular biology experiments. Unfortunately, the intrinsic loss in accuracy of the latter approaches imposes significant compromises. Therefore, devoting continuous efforts to increase the accuracy of existing CG force fields remains an important task. In this contribution we presented a significant update of the SIRAH force field for proteins. As stated in the title, improvements can be classified in three categories:

***Altius***: the newly introduced implementation in AMBER facilitates its use reaching a higher number of potential users. Owing to its straightforward implementation and use of a classical Hamiltonian takes profit of all possible MD implementations in GROMACS and AMBER. Along the same line, the SIRAH Tools enlarged the spectra of mapping definitions, which facilitate the set up of CG simulations.

Although a universal recipe for parameter development is not available on SIRAH, the physicochemical vision of beads based on functional groups provided in Figure 2 will facilitate the development of a higher number of molecules, enriching the biological diversity amenable to be studied with SIRAH. Indeed, parameters for CG glycans, metallic ions and different phospholipidss compatible with this protein force field are currently being tested. These sets of new interaction parameters will offer in the near future the possibility to address highly complex systems containing glycosylated and metallo-proteins, polysaccharides, membranes and nucleic acids.

***Fortius***: the robustness of SIRAH 2.0 resulted not only in a more accurate description of a set of structures but also a smaller dispersion in relation to the experimental data, reaching in some cases nearly atomistic resolution. On the other hand, the CaM system used as test case revealed several interesting features that strongly suggest the force field could achieve at least a very good qualitative description. The simulation of helical peptides showed the expected sequence dependent conformational behavior. Moreover, the CaM binding peptide unfolded only when isolated, evidencing also sensitivity to the quaternary context. The fully unbiased wrapping of EF-hands onto the binding peptide underlines this attribute. Under this perspective, SIRAH 2.0 can be considered a significant upgrade that comes at no increase of computational cost, as the functional form of the Hamiltonian, the number of beads in each moiety, and their topologies remained the same.

***Citius***: the straightforward implementation of the new version of the force field fully profits from all available options present in GROMACS and AMBER packages. In particular the GPU implementations making possible the simulation of middle sized systems at a rate of few μs per day in GPU-accelerated desktop computers.

Finally, it is important to recall that CG models in general, and ours in particular, are not to be considered as a universal strategy for reducing computational cost. Although SIRAH was developed as a generic force field for biological systems, there will always be problems not amenable to be solved by coarse-graining. Those comprise systems in which is necessary to consider atomistic details, H-bonds or single water molecules, but also others in which the intrinsic reduction of degrees of freedoms precludes the unbiased representation of particular conformations. It is important for computational biologists to keep in mind such limitations and choose the most appropriated technique to solve their problems of interest.

## ACKNOWLEDGMENTS

This work partially funded by FOCEM (MERCOSUR Structural Convergence Fund), COF 03/11. M.R.M. and S.P. belong to the SNI program of ANII. M.S. and F.K. were partially financed by postgraduate fellowships by ANII. E.E.B. is beneficiary of a postdoctoral fellowship of CONICET (Consejo Nacional de Investigaciones Científicas y Técnicas, Argentina). Graphic cards Tesla K40c and Titan-Xp, used in this research were donated by the NVIDIA Corporation.

## SUPPLEMENTARY MATERIAL

**Table ST1.** Detailed results on the benchmark data set simulated with SIRAH 2.0. Average values and standard deviations were calculated on triplicates. Regular rows correspond to GROMACS, while yellow rows correspond to AMBER.

| PDB | N | RMSD (nm) | Rgyr (nm) | SAS (nm<sup>2</sup>) |  | Secondary structure |  |  | Contacts |
| --- | --- | --- | --- | --- | --- | --- | --- | --- | --- |
|  |  |  |  | Hydrophobic | Hydrophilic | Helix | Extended | Coiled | Conservation / Accuracy |
| 1TQG | 105 | 0.23 (0.06) | [1.38] 1.37 (0.02) | [6.7] 7.67 (0.76); [58.8] 63.67 (3.19) |  | [86.7] 80.50 (2.81); [1.9] 3.2 (0.53); [11.4] 16.30 (3.28) |  |  | 89.90 (2.35) / 90.63 (2.47) |
|  |  | 0.29 (0.01) | 1.46 (0.03) | 8.33 (1.11); | 62.70 (0.85) | 85.03 (1.48); | 0.13 (0.12); | 14.80 (1.64) | 88.20 (0.87) / 89.73 (1.33) |
| 1R69 | 63 | 0.65 (0.09) | [1.01] 1.12 (0.05) | [3.2] 8.90(1.42); [38.2] 44.20 (3.32) |  | [69.8] 60.33 (2.62); [0] 0.63 (0.76); [30.2] 39.03 (2.14) |  |  | 78.80 (1.55) / 80.20 (3.98) |
|  |  | 0.43 (0.04) | 1.13 (0.03) | 7.23 (1.53); | 44.50 (2.26) | 60.80 (2.79); | 1.43 (1.88); | 37.73 (1.27) | 81.40 (3.08) / 81.50 (4.24) |
| 2CKX | 83 | 0.42 (0.03) | [1.17] 1.28 (0.03) | [7.7] 8.73 (0.21); [45.5] 57.27 (0.06) |  | [66.3] 64.10 (1.47); [6] 7.87 (0.64); [27.7] 28.07 (0.81) |  |  | 83.37 (0.23) / 85.80 (1.13) |
|  |  | 0.38 (0.04) | 1.28 (0.05) | 9.87 (1.01); | 55.33 (3.09) | 63.20 (1.00); | 4.73 (1.48); | 32.10 (1.13) | 83.63 (1.15) / 87.57 (0.68) |
| 1BKR | 108 | 0.60 (0.09) | [1.24] 1.28 (0.05) | [6] 10.00 (0.26); [57.8] 65.10 (2.60) |  | [58.3] 49.33 (1.10); [3.7] 10.80 (1.73); [38] 39.93 (2.40) |  |  | 75.67 (0.68) / 78.73 (2.05) |
|  |  | 0.32 (0.02) | 1.30 (0.02) | 11.93 (1.45); | 60.80 (1.60) | 54.30 (1.05); | 6.33 (2.36); | 39.37 (2.71) | 80.57 (0.80) / 82.27 (2.00) |
| 1RA4 | 117 | 0.38 (0.03) | [1.23] 1.21 (0.03) | [6.5] 7.50 (0.40); [59.7] 66.27 (2.59) |  | [48.7] 39.43 (3.87); [19.7] 13.27 (0.38); [31.6] 47.30 (3.80) |  |  | 76.23 (1.80) / 76.47 (3.23) |
|  |  | 0.28 (0.02) | 1.25 (0.02) | 7.40 (1.15); | 63.53 (0.76) | 44.93 (0.78); | 16.67 (1.07); | 38.40 (1.80) | 83.53 (1.52) / 81.97 (0.29) |
| 2KYR | 108 | 0.39 (0.05) | [1.27] 1.25 (0.02) | [7.7] 9.87(0.12); [57.3] 61.27 (0.81) |  | [46.3] 40.80 (2.95); [32.4] 23.87 (1.19); [21.3] 35.33 (1.76) |  |  | 76.17 (2.42) / 84.13 (2.00) |
|  |  | 0.32 (0.08) | 1.30 (0.00) | 8.83 (0.59); | 59.90 (1.82) | 44.30 (1.80); | 24.90 (2.82); | 30.73 (4.57) | 80.77 (2.91) / 85.60 (1.23) |
| 2VIM | 104 | 0.44 (0.08) | [1.22] 1.23 (0.02) | [5.4] 7.40 (1.30); [52.2] 61.60 (0.30) |  | [46.2] 39.50 (0.44); [36.5] 20.20 (2.00); [17.3] 40.33 (1.72) |  |  | 76.63 (0.78) / 81.47 (2.49) |
|  |  | 0.24 (0.02) | 1.27 (0.02) | 6.43 (0.12); | 59.97 (1.65) | 39.53 (1.10); | 28.53 (1.11); | 31.97 (2.08) | 84.43 (1.89) / 88.03 (0.06) |
| 1CRN | 46 | 0.21 (0.05) | [0.97] 0.97 (0.04) | [7.6] 9.23 (0.12); [24.4] 27.07 (1.20) |  | [45.7] 41.83 (2.47); [23.9] 12.60 (3.01); [30.4] 45.60 (5.20) |  |  | 88.47 (2.68) / 96.93 (0.35) |
|  |  | 0.25 (0.08) | 1.00 (0.04) | 9.63 (0.29); | 26.83 (2.01) | 43.93 (1.19); | 10.27 (3.75); | 45.87 (3.13) | 88.80 (1.05) / 95.77 (0.55) |
| 1ORC | 64 | 0.49 (0.05) | [1.08] 1.09 (0.04) | [4.9] 5.97 (0.76); [43.9] 47.77 (1.33) |  | [40.6] 39.43 (0.84); [35.9] 19.33 (1.37); [23.4] 41.20 (2.16) |  |  | 86.23 (2.78) / 84.47 (2.25) |
|  |  | 0.38 (0.07) | 1.16 (0.06) | 6.63 (1.67); | 47.60 (2.61) | 38.17 (1.79); | 18.70 (2.82); | 43.13 (4.54) | 85.33 (3.12) / 85.37 (2.37) |
| 1PGB | 56 | 0.36 (0.02) | [1.03] 1.01 (0.01) | [2.7] 3.07 (0.40); [37.7] 42.57 (1.53) |  | [26.8] 23.63 (3.84); [51.8] 31.27 (6.08); [21.4] 45.10 (3.20) |  |  | 88.70 (0.96) / 84.60 (1.25) |
|  |  | 0.32 (0.07) | 1.08 (0.03) | 4.00 (0.66); | 43.73 (1.44) | 24.93 (0.06); | 29.07 (7.70); | 46.03 (7.70) | 88.53 (1.60) / 88.80 (6.08) |
| 2IUG | 110 | 0.65 (0.08) | [1.28] 1.43 (0.03) | [7.7] 9.23 (1.16); [56.5] 70.30 (1.87) |  | [22.7] 20.77 (1.33); [37.3] 20.43 (1.97); [40] 58.90 (1.61) |  |  | 70.63 (3.54) / 80.47 (2.44) |
|  |  | 0.48 (0.10) | 1.39 (0.02) | 8.93 (1.00); | 68.23 (1.22) | 19.83 (0.99); | 26.27 (2.39); | 53.90 (2.42) | 78.87 (2.93) / 85.67 (3.44) |
| 1QYO<sup>a</sup> | 236 | 0.40 (0.04) | [1.75] 1.83 (0.05) | [8.1] 12.03 (1.10); [104.9] 112.93 (3.89) |  | [7.2] 6.83 (1.06); [55.5] 47.50 (1.47); [37.3] 45.63 (1.62) |  |  | 76.90 (2.43) / 79.83 (1.82) |
|  |  | 0.31 (0.04) | 1.81 (0.02) | 12.80 (1.47); | 114.20 (1.44) | 8.47 (1.42); | 47.43 (1.01); | 44.07 (1.24) | 80.60 (0.78) / 83.90 (0.72) |
| 1OPS | 64 | 0.52 (0.19) | [1.01] 1.02 (0.02) | [8.6] 9.20 (0.79); [31.1] 37.17 (0.47) |  | [4.7] 1.60 (1.42); [56.2] 27.90 (5.72); [39.1] 70.47 (4.30) |  |  | 63.43 (2.78) / 73.43 (5.36) |
|  |  | 0.41 (0.08) | 1.05 (0.03) | 10.00 (1.14); | 36.73 (2.25) | 2.37 (0.76); | 31.03 (3.84); | 66.60 (4.08) | 71.90 (2.69) / 79.87 (1.36) |
| 1GYV | 120 | 0.30 (0.01) | [1.44] 1.48 (0.01) | [8.6] 10.53 (0.61); [60.4] 63.97 (1.05) |  | [1.7] 2.47 (1.44); [61.7] 47.77 (2.24); [36.7] 49.77 (1.80) |  |  | 81.50 (1.31) / 85.93 (3.00) |
|  |  | 0.29 (0.05) | 1.49 (0.01) | 11.17 (1.45); | 66.33 (1.10) | 0.83 (0.06); | 47.97 (1.90); | 51.20 (1.85) | 80.77 (2.22) / 86.27 (2.29) |
| 2E3H | 76 | 0.38 (0.07) | [1.12] 1.18 (0.03) | [7.6] 8.70 (1.01); [43.4] 51.03 (1.57) |  | [0] 0.00 (0.00); [48.7] 30.93 (2.04); [51.3] 69.03 (2.05) |  |  | 71.37 (3.77) / 78.23 (5.61) |
|  |  | 0.39 (0.15) | 1.17 (0.02) | 9.00 (0.35); | 50.67 (0.32) | 0.00 (0.00); | 36.90 (4.17); | 63.10 (4.17) | 76.70 (4.23) / 77.77 (3.80) |
| 1PWT | 61 | 0.37 (0.05) | [1.03] 1.07 (0.03) | [5.4] 6.50 (0.78); [43.5] 47.77 (3.31) |  | [0] 0.00 (0.00); [60.7] 35.10 (5.99); [39.3] 64.90 (5.99) |  |  | 73.17 (3.90) / 87.67 (1.72) |
|  |  | 0.25 (0.06) | 1.15 (0.04) | 7.40 (0.35); | 48.50 (2.05) | 0.00 (0.00); | 41.90 (8.89); | 58.10 (8.89) | 82.97 (4.39) / 91.30(1.95) |
| 2O31 | 67 | 0.47 (0.12) | [1.05] 1.23 (0.10) | [9.5] 12.90 (1.04); [39.8] 46.93 (1.97) |  | [0] 0.27 (0.31); [62.7] 38.30 (6.97); [37.3] 61.40 (6.88) |  |  | 75.47 (3.93) / 80.30 (8.98) |
|  |  | 0.32 (0.08) | 1.16 (0.05) | 10.43 (0.06); | 44.97 (1.16) | 0.03 (0.06); | 42.00 (6.05); | 57.97 (5.99) | 79.77 (6.31) / 89.47 (2.57) |
| 1XX8 | 66 | 0.42 (0.13) | [1.12] 1.17 (0.06) | [3.5] 3.30 (0.78); [50] 57.13 (2.59) |  | [16.7] 12.37 (1.54); [42.4] 33.13 (2.29); [40.9] 54.50 (1.97) |  |  | 81.13 (7.25) / 85.53 (5.06) |
|  |  | 0.30 (0.01) | 1.19 (0.01) | 3.47 (1.00); | 59.90 (1.85) | 12.67 (0.12); | 33.57 (3.65); | 53.77 (3.55) | 87.60 (1.00) / 87.93 (3.18) |

<sup>a</sup> The buried residue E222 was modeled as a protonated glutamate as it was predicted to be in that state by the PDB2PQR server.

**Figure S1.**
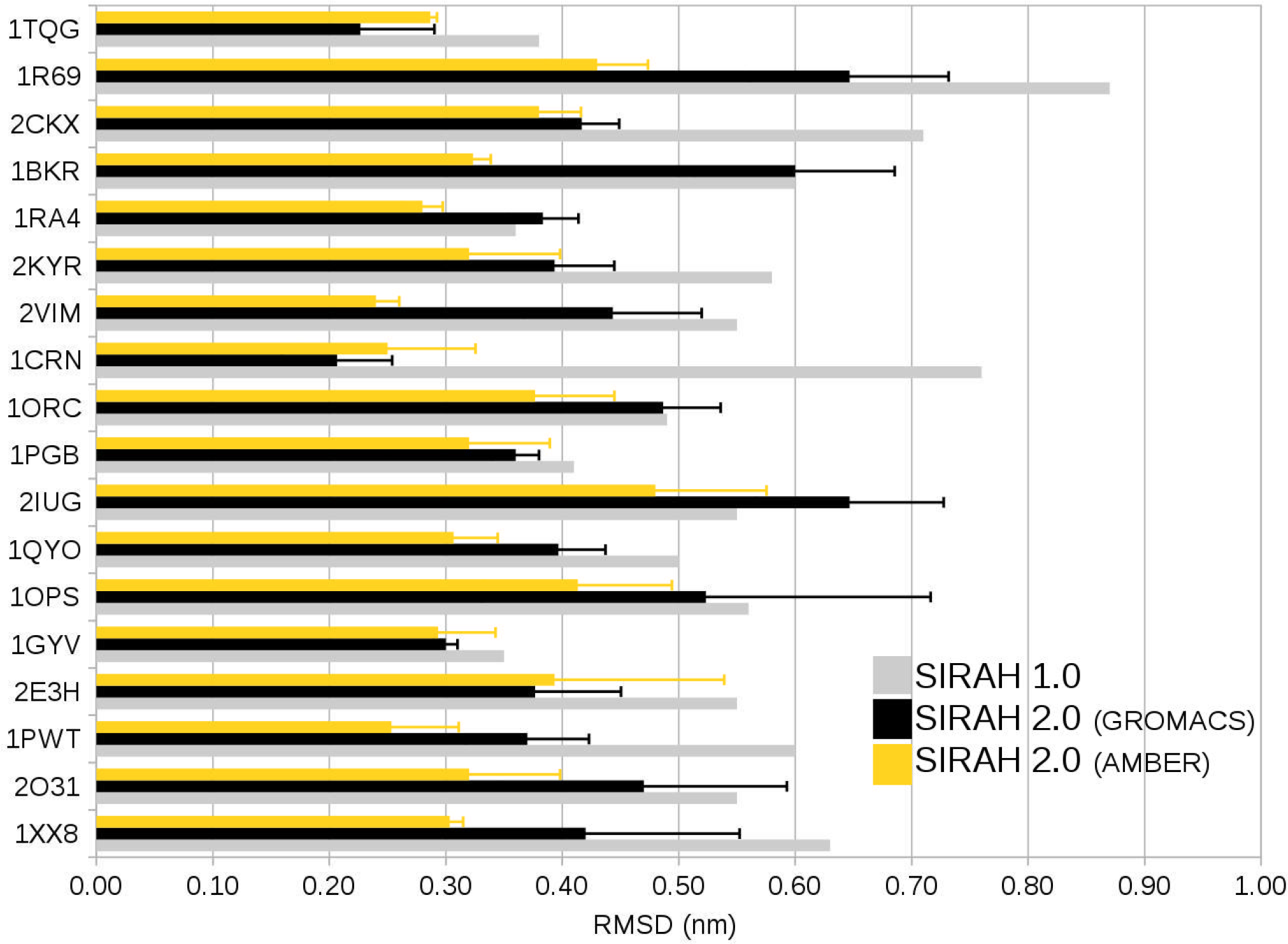
Graphical representation of RMSD results for each protein in the benchmark data set. Average values and standard deviations were calculated on triplicates.

